# Reciprocal regulation of Shh trafficking and H_2_O_2_ levels via a noncanonical BOC-Rac1 pathway

**DOI:** 10.1101/2021.07.15.452471

**Authors:** Marion Thauvin, Irène Amblard, Christine Rampon, Aurélien Mourton, Isabelle Queguiner, Chenge Li, Arnaud Gautier, Alain Joliot, Michel Volovitch, Sophie Vriz

## Abstract

Among molecules that bridge environment, cell metabolism, and cell signaling, H_2_O_2_ recently appeared as an emerging but central player. Its level depends on cell metabolism and environment and was recently shown to play key roles during embryogenesis, contrasting with its long-established role in disease progression. We decided to explore whether the secreted morphogen Sonic hedgehog (Shh), known to be essential in a variety of biological processes ranging from embryonic development to adult tissue homeostasis and cancers, was part of these interactions. Here, we report that H_2_O_2_ levels control key steps of Shh delivery and that physiological *in vivo* modulation of H_2_O_2_ levels changes Shh distribution and tissue patterning. A feedback loop exists in which Shh trafficking controls H_2_O_2_ synthesis via a non-canonical BOC-Rac1 pathway, leading to cytoneme growth. Our findings reveal that Shh directly impacts its own distribution, thus providing a molecular explanation for the robustness of morphogenesis to both environmental insults and individual variability.

## Introduction

Hedgehog proteins (Hh in *Invertebrates* and Shh or paralogs in *Vertebrates*, collectively Hhs) are secreted morphogens playing key roles in biological processes ranging from embryonic development, proliferation, adult tissue homeostasis, and cancers (Briscoe and Therond, 2013; Ingham and McMahon, 2001; McMahon et al., 2003). Signaling by Hhs implies a complex protein journey that includes several steps incompletely characterized, and possibly depending on physiological context (Guerrero and Kornberg, 2014; Hall et al., 2019). Recently, we observed that Shh compartimentation was modified *ex vivo* by H_2_O_2_ (Gauron et al., 2016), suggesting that H_2_O_2_ could regulate Shh secretion process. Besides, preliminary data suggested that Shh controls H_2_O_2_ levels during cell plasticity and tissue remodeling in adults (Meda et al., 2016). Hydrogen peroxide (H_2_O_2_) has long been exclusively considered as a deleterious molecule damaging cellular integrity and function. It is now becoming evident that H_2_O_2_ also contributes to *bona fide* physiological processes notably through protein cysteine targeting (Holmstrom and Finkel, 2014; Sies, 2017; Sies and Jones, 2020).

The physiological role of H_2_O_2_ in living systems has gained increased interest and has been investigated in different models of development and regeneration (Breus and Dickmeis, 2021; Coffman and Su, 2019; Rampon et al., 2018; Timme-Laragy et al., 2018). Several studies have revealed strong spatio-temporal variation of H_2_O_2_ levels in live animals of various species (fly, nematode, zebrafish, *Xenopus*) during these processes (Albadri et al., 2019; Albrecht et al., 2011; Amblard et al., 2020b; Bazopoulou et al., 2019; Gauron et al., 2016; Gauron et al., 2013; Han et al., 2018; Katikaneni et al., 2020; Knoefler et al., 2012; Love et al., 2013; Meda et al., 2016; Mendieta-Serrano et al., 2019; Niethammer et al., 2009; Pak et al., 2020; Tao et al., 2017). The correlation between mild oxidative bursts and developmental events led to early suggestions of a mechanistic link between them (Covarrubias et al., 2008; Hernandez-Garcia et al., 2010; Sies, 2014). Since the patterning of a developing embryo relies on the graded activity of morphogens (Lander, 2007; Rogers and Schier, 2011; Turing, 1952), we decided to precisely determine by which mechanism H_2_O_2_ could regulate Shh trafficking. We set up a quantitative assay to measure the efficiency of each step of Shh’s journey and demonstrate that H_2_O_2_ inhibits Shh secretion but enhances Shh internalization. Shh internalization per se enhances endogenous H_2_O_2_ levels via a Rac1/NADPH oxidase pathway that induces filopodia growth and regulates Shh trafficking in an H_2_O_2_/Shh feedback loop.

## Results and Discussion

### *H_2_O_2_ affects Shh trafficking* ex vivo

To analyse the mechanism by which H_2_O_2_ levels could affect Shh trafficking, we used an *ex vivo* system enabling accurate quantification of this process. HeLa cells were chosen to avoid complex feedback loops: these cells do not express endogenous Shh or respond to the canonical Shh signaling pathway but otherwise express pathway components known to be involved in Shh trafficking (lipidation, access to the extracellular space, endocytosis, and delivery to receiving cells, Supplementary Table S1). Shh trafficking from producing to receiving cells is a circuitous journey. While some aspects are still a matter of debate, there is no doubt that cis-endocytosis in producing cells can be part of the process (Guerrero and Kornberg, 2014; Petrov et al., 2017). Making use of Shh constructs (mouse sequence) tagged according to (Chamberlain et al., 2008) to preserve the qualitative properties of the protein, we first verified that HeLa cells recapitulated the overall traffic of Shh (Fig. 1A-D). To distinguish different steps of the Shh journey (Fig. 1A), we exploited a double tag combining a fluorogen-activating peptide (YFAST) and a classical fluorescent protein (mCherry). Contrary to mCherry, which chromophore takes time to mature, YFAST fluoresces instantly after fluorogen addition (Plamont et al., 2016)’ allowing Shh detection from the beginning of its journey in the ER (step 1 in Fig. 1A) too early for mCherry detection. However, YFAST is sensitive to pH and cannot fluoresce in endosomes (step 2 in Fig. 1A), contrary to mCherry. When finally sent for receiving cells (step 3 in Fig. 1A) Shh should be detectable via both tags. Transfection of HeLa cells with such a construct indeed confirmed the usefulness of this cell line for our purposes (Fig. 1B). At a steady state, the majority of cells display diffuse green staining, including the ER, as well as large red spots about the size of endosomes (Fig. 1B). HeLa cells with cellular protrusions associated with yellow puncta were found, indicating detection via both tags of Shh en route to receiving cells (Fig. 1B). The end of the journey was also easy to image by co-culturing transfected and untransfected HeLa cells. As shown in Fig. 1C, when transfected and untransfected cells were nearby, Shh-mCherry could be detected in the untransfected cells, and the arrangement of cytoplasmic protrusions between the two cells suggested a transfer occurring via filopodia.

**Fig. 1.**
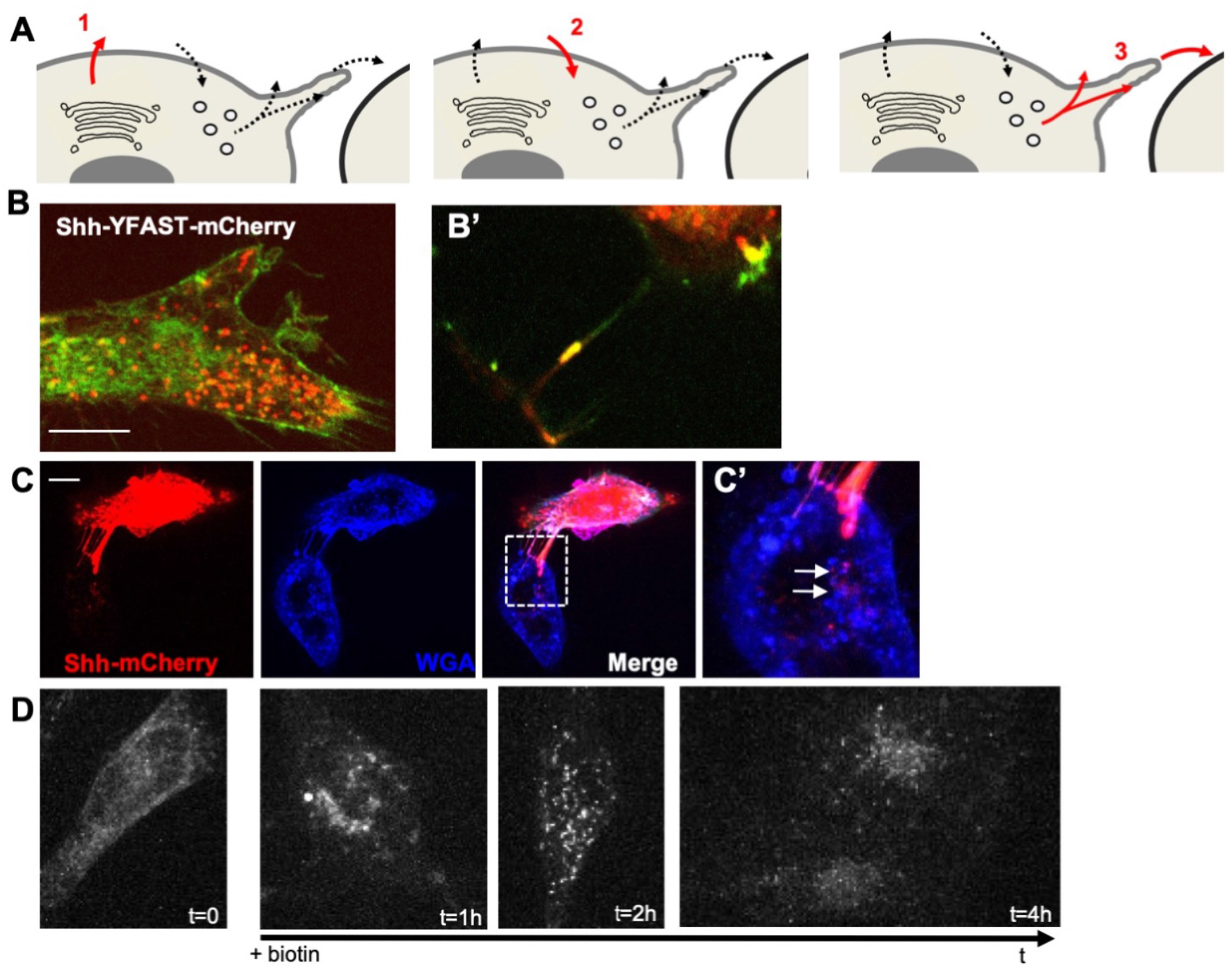
HeLa cells recapitulate typical events of Shh trafficking *ex vivo*. (**A**) Schematic view of important steps analysed in HeLa cells: 1) first secretion; 2) endocytosis; 3) dispatching to receiving cells. (**B**) Double tagging with YFAST and mCherry allows the detection of Shh in three different compartments. Left panel: in a steady state, a cell producing Shh-YFAST-mCherry exhibits both a diffuse green signal (YFAST detected early in the endoplasmic reticulum, when mCherry has not yet matured) and an abundant red vesicular signal (mCherry detected in the endosomes, where the pH prevents YFAST detection) and in filopodia where both tags are detected (**B’**). (**C**) In co-culture, pairs of Shh-producing (mCherry signal) and non-producing cells often display filopodia between them. Large red dots in the non-producing cell indicate Shh transfer to recipient cells, and their position suggests delivery via the filopodia (white arrows in the enlarged inset in C’). (**D**) Shh-Sbp-mCherry secretion synchronized with the RUSH system displays classic timing: Shh is correctly hooked in the endoplasmic reticulum before biotin addition, reaches the Golgi by 30-40 min after biotin addition, is secreted at approximately 1 h, can be easily detected within endosomes of the producing cell at 2 h, and can be visualized in non-producing cells at 4 h. Scale bars, 10 μm.

To reach an adequate level of precision in the analysis of H_2_O_2_ effect on Shh trafficking, we needed to synchronize its secretion. We thus made use of the RUSH system (Boncompain et al., 2012), where Shh fused to the streptavidin binding peptide (SBP) is retained in the ER of cells expressing streptavidin fused to the KDEL retrieving signal until the addition of biotin. As shown in Fig. 1D, before biotin addition (t=0), Shh-SBP-AFP (mCherry here) was hooked in the ER, and biotin addition allowed us to calibrate the timing of the Shh journey using realtime imaging. It takes approximately 30 to 40 min for the bulk of Shh to reach the Golgi (very similar to many secreted proteins in these conditions, e.g., Wnt3a (Moti et al., 2019)), secretion is abundant from 1 h, localization in endosomes is conspicuous between 2 and 3 h, and detection in (or at the surface of) non-producing cells lasts up to 5 h and then disappears. HeLa cells thus display typical features of Shh trafficking and can be used to analyse the potential effects of modifying H_2_O_2_ levels if we have means to rigorously quantify Shh in different compartments at different steps of the process.

In addition, we recently developed (Amblard et al., 2020a) quantitative assays to track the different steps of a protein journey by combining the RUSH (Boncompain et al., 2012) and HiBit systems (Dixon et al., 2016) (HiBit assay, Promega) (Fig. 2). The Hibit system is constituted by a split luciferase: when in the same compartment, the two luciferase moieties, HiBiT (Sbi in plasmid names) and LgBiT (G/Lbi in plasmid names) may spontaneously assemble and restore the luciferase activity that can be measured by substrate addition. These assays are inducible, quantitative, and specifically adapted to protein trafficking. We then set up this quantitative assay to analyse separately each step of Shh’s journey. This strategy allowed us to determine the optimal time frame for analysing each step of Shh trafficking in cell culture, i.e. secretion, endocytosis, and delivery *via* filopodia. These quantitative results are consistent with our direct fluorescence microscopy observations (Fig. 1D) and were combined with redox modulation tools to test the hypothesis of the redox regulation of Shh trafficking.

**Fig. 2.**
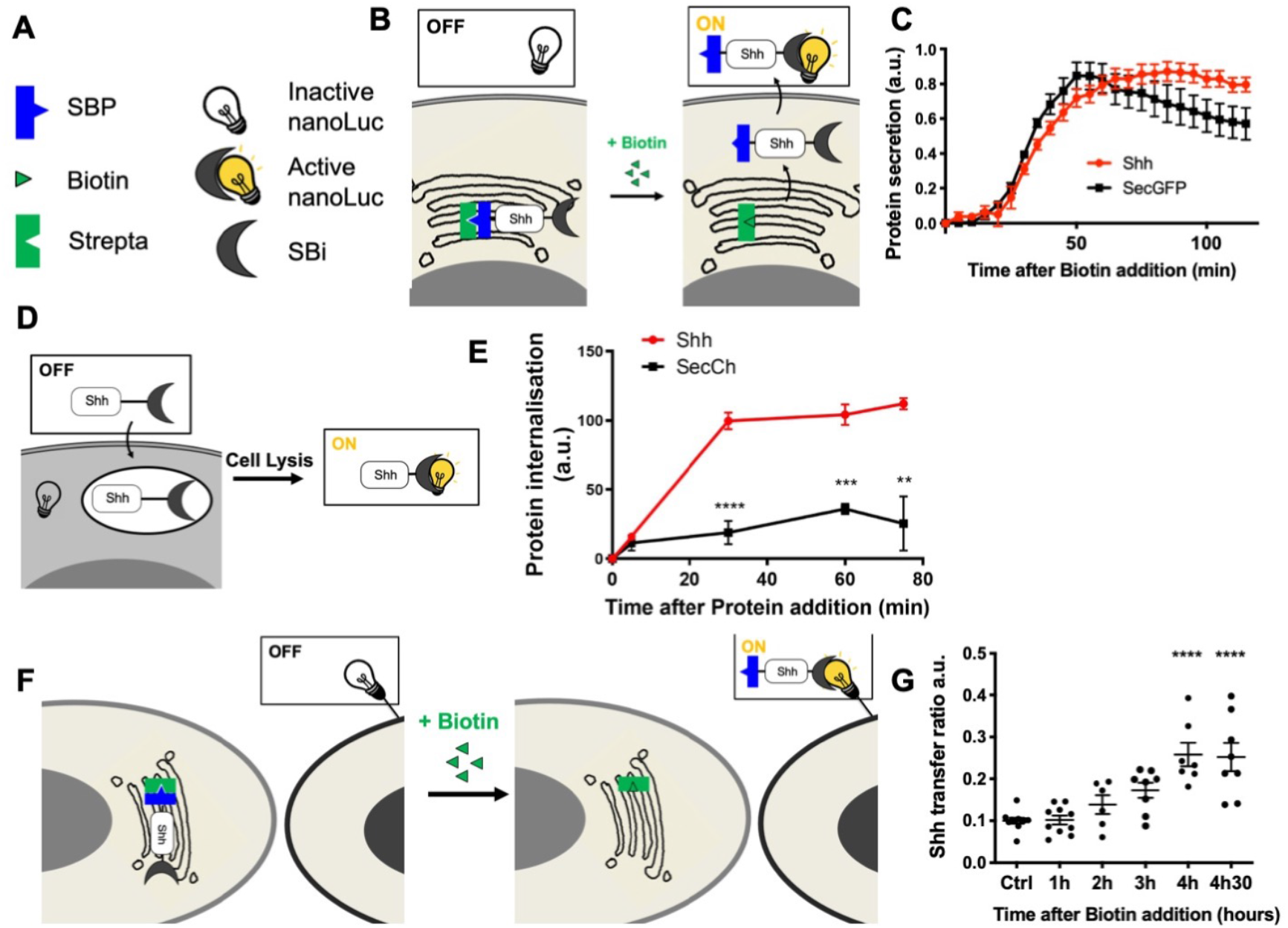
Quantitative monitoring of the journey of Shh *ex vivo*. (**A-C**) Quantifying Shh secretion by HeLa cells. (**A, B**) Schematic representation of the tools (**A**) and assay (**B**). (**A**) Strepta: core streptavidin linked to the KDEL hook; SBP: streptavidin binding protein domain linked to Shh or secGFP; SBi: small fragment of split nanoluciferase (HiBiT) linked to Shh or secGFP; inactive nanoLuc: large fragment of split nanoluciferase (LgBiT); active nanoLuc: reconstituted nanoluciferase. (**B**) Synchronized HeLa cells constitutively express the hook, and they express Shh fused to SBP and HiBiT (Shh-SBP-HiBiT) under doxycycline control. Purified LgBit added to the medium binds HiBiT when Shh reaches the extracellular space, and nanoLuc activity gives a measure of secretion. (**C**) Time course of Shh synchronized secretion compared to the control (secGFP fused to SBP and HiBiT). (**D-E**) Quantitating Shh internalization by HeLa cells. (**D**) Schematic representation of the assay. Shh fused to HiBiT (Shh-HiBiT), purified from the conditioned medium of producing HeLa cells and calibrated *in vitro*, was added to HeLa cells expressing inactive nanoLuc (LgBiT) in the cytosol. Upon cell lysis, nanoluciferase is reconstituted from endocytosed Shh-HiBiT and cytoplasmic LgBiT, allowing quantification of internalization. (**E**) Time course of Shh internalization compared to the control (secCherry fused to HiBiTI, prepared in parallel to Shh). (**F-G**) Quantitating Shh delivery to recipient HeLa cells. (**F**) Schematic representation of the assay. Two HeLa cell populations are co-cultured. In the first one, synchronization of Shh release is achieved as in the secretion assay but for a longer period of time. The second population expresses inactive nanoLuc (LgBiT) at the cell surface (anchored via the CD4 transmembrane domain). Active nanoluciferase is reconstituted by the transport of tagged Shh (Shh-SBP-HiBiT) from a donor cell to the surface of a receiving cell. (**G**) Time-course analysis of Shh delivery to recipient cells becomes strongly significant approximately 3 h after biotin addition. Ctrl: without biotin. Details on statistics in Methods.

First, we studied the effects of H_2_O_2_ on primary Shh secretion. To increase H_2_O_2_ levels, we expressed inducible D-aminoacid oxidase (DAO), which produces H_2_O_2_ in the presence of D-Ala and is not expressed in HeLa cells (Haskew-Layton et al., 2010; Matlashov et al., 2014). For cells expressing DAO and treated with 10 mM D-Ala, we observed a specific reduction in Shh secretion, not observed with a secGFP construct taken as a control (Fig. 3A-B). This result is in close agreement with our previous observations: after oxidative treatment, cells showed reduced Shh secretion, and a pool of Shh was trapped in the Golgi apparatus (Gauron et al., 2016). Conversely, reduction of H_2_O_2_ levels by the direct addition of Catalase (CAT) in cell culture (Amblard et al., 2020b), enhanced Shh but not SecGFP secretion (Fig. 3A,C). Next, we applied the same treatments to study Shh endocytosis on cells incubated with conditioned media of Shh expressing cells. Compared to control conditions (secreted mCherry conditioned media), enhancing H_2_O_2_ levels with DAO (by addition of D-Ala) stimulated Shh endocytosis (Fig. 3D-E). Conversely, the reduction in H_2_O_2_ levels with CAT treatment reduced Shh endocytosis (Fig. 3D,F). Finally, we studied the effects of H_2_O_2_ level modulation on Shh delivery to recipient cells in the co-culture assay. Cells expressing DAO and treated with D-Ala (but not untreated cells) showed increased Shh delivery to recipient cells (Fig. 3G-H), while a reduction in H_2_O_2_ levels with extracellular catalase had the opposite effect (Fig. 3G,I). Altogether, these results indicate that physiological variations in H_2_O_2_ levels impact Shh trafficking *ex vivo*. This raises the interesting possibility that the heterogenous H_2_O_2_ distribution could polarize Shh secretion and endocytosis *in vivo*.

**Fig. 3:**
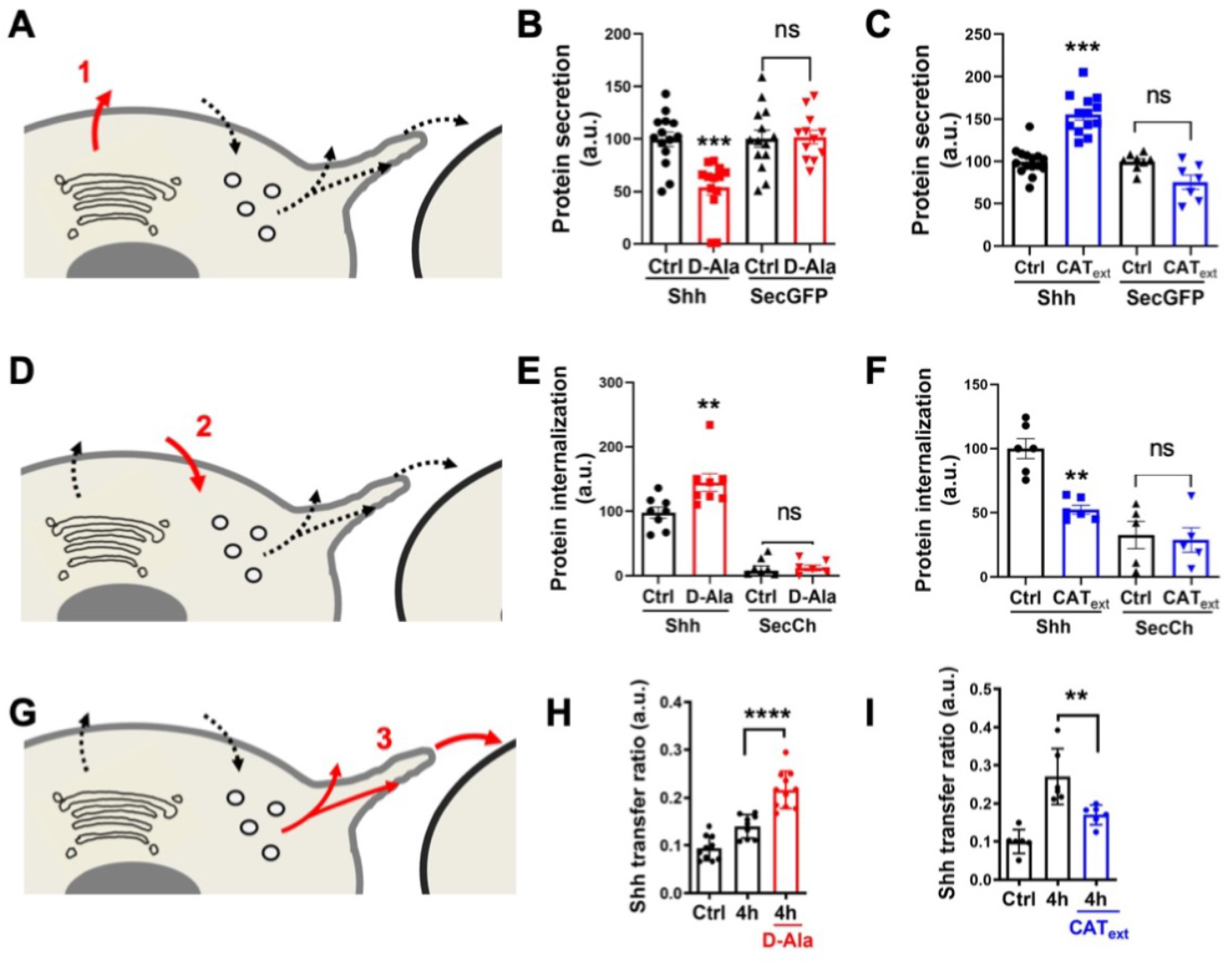
H_2_O_2_ affects Shh secretion, endocytosis and delivery in HeLa cells. (**A, D, G**) Schematic steps (Fig. 3): secretion (**A**), endocytosis (**D**), delivery (**G**). **A-C**, Effects of increased (**B**) or decreased (**C**) H_2_O_2_ levels on secretion of Shh or secGFP from cells expressing Lck-DAO supplemented (or not) with D-Ala (**B**) or cells treated (or not) with extracellular catalase (**C**). (**D-F**) Effects of increased (**E**) or decreased (**F**) H_2_O_2_ levels on Shh or secCherry endocytosis by cells expressing Lck-DAO supplemented (or not) with D-Ala (**E**) or cells treated (or not) with extracellular catalase (**F**). **G-I**, Effects of increased (**H**) or decreased (**I**) H_2_O_2_ levels on delivery to recipient cell surface of cells expressing Lck-DAO supplemented (or not) with D-Ala (**H**) or cells treated (or not) with extracellular catalase (**I**). Details on statistics in Methods.

### H_2_O_2_ levels are dynamic in time and space in the embryonic spinal cord

Between 25 and 45 h post fertilization (hpf) in zebrafish, Shh morphogen is first expressed in the medial floor plate (MFP) consisting of a single row of cells at the central midline, then extends to the flanking lateral floor plate (LFP), playing an important role in the neuroglial switch (Danesin and Soula, 2017). Using the improved H_2_O_2_ ratiometric probe HyPer7 (Pak et al., 2020), we first measured H_2_O_2_ levels in Shh-expressing cells in the spinal cord of live zebrafish embryos during this time window (Fig. 4). Quantitative analysis of the H_2_O_2_ signal demonstrated a regular and significant decrease (approximately 25%) in H_2_O_2_ levels in the MFP between 25 and 45 hpf (Fig. 4A-B). A close-up view of the MFP showed that, in addition to an overall decrease in concentration, the spatial distribution of H_2_O_2_ levels varies over time (Fig. 4C-D). Time-lapse analysis of these cells revealed a dynamic intracellular distribution of the HyPer7 signal between 25 and 45 hpf. At the beginning and end of this period, H_2_O_2_ levels are homogeneously spread throughout the cell. In between, however, a distinct gradient is transiently established from higher concentrations apically to lower concentrations basolaterally, and most marked at 31 hpf (Fig. 4C-D). Thus, during this neurogenesis period, when Shh induces oligodendrocyte precursor cells (OPCs), H_2_O_2_ levels decrease over time and exhibit a transiently polarized distribution within MFP cells.

**Fig. 4:**
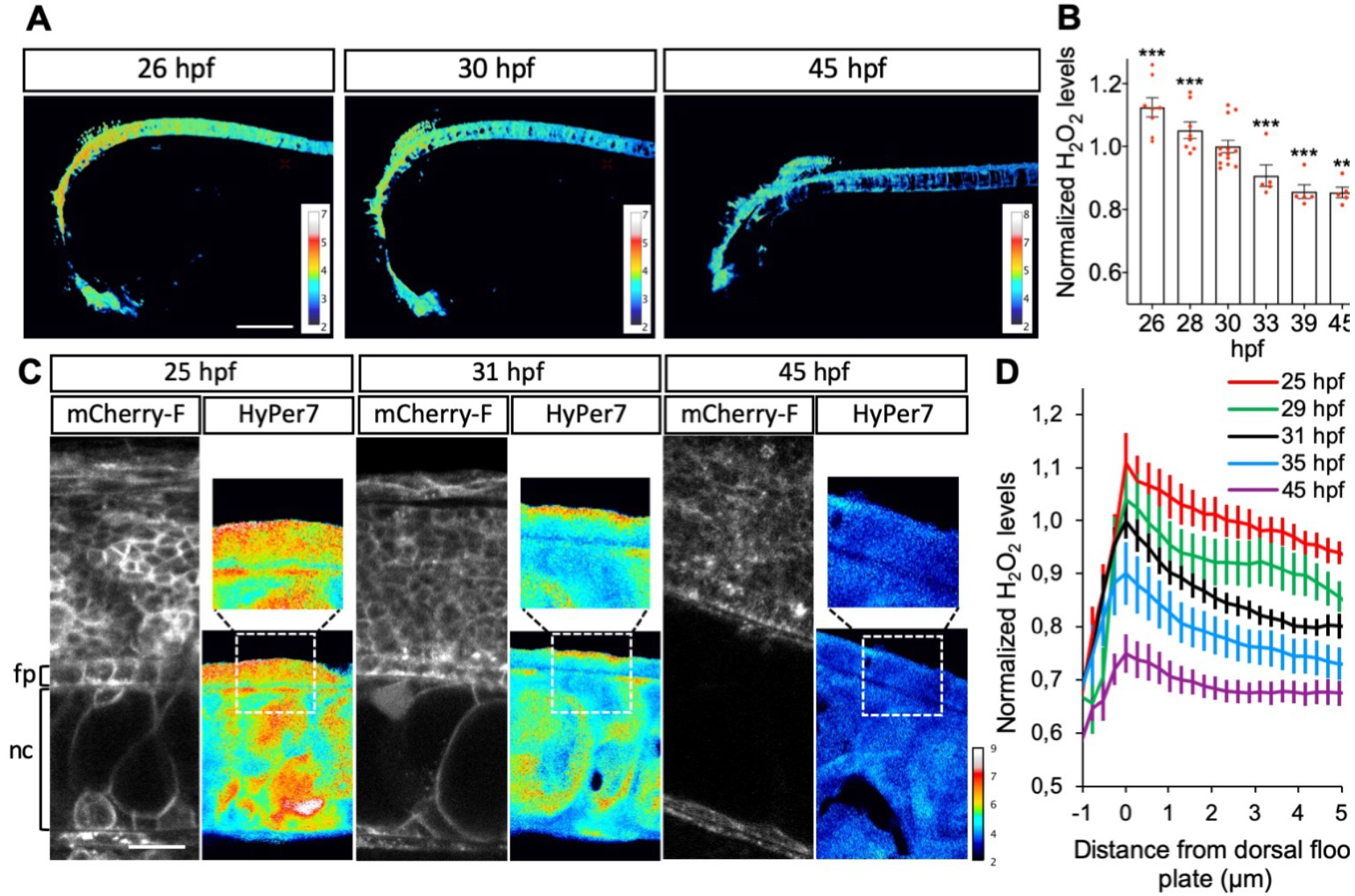
H_2_O_2_ levels are dynamic in time and space in the floor plate. (**A-B**) Variation in time. (**A**) H_2_O_2_ levels in Tg(2.4Shha-ABC:Gal4-FF; UAS:Hyper7) zebrafish transgenic embryos at different times after fertilization as indicated (lateral view, anterior to the left). Scale bar, 200 μm. (**B**) Variations in H_2_O_2_ levels at different stages. Statistics are presented compared to 30 hpf. (**C-D**) Transient dorso-ventral gradient. (**C**) Embryos injected at the one-cell stage with mRNA for farnesylated mCherry were imaged for membranes (left panels: mCherry-F) and H_2_O_2_ (right panels: HyPer7; bottom: same field as in the left panel; top: enlarged view as indicated by the dotted line). fp: floor plate; nc: notochord. Scale bar: 20 μm. (**D**) Variations in H_2_O_2_ levels along the apico-basal axis of MFP cells. Details on statistics in Methods.

### H_2_O_2_ impacts filopodial formation and Shh cellular targets in the embryonic spinal cord

Shh can be delivered to the receiving cells via filopodia, and it has been proposed that the length of filopodia impacts Shh distribution (Fairchild and Barna, 2014; Gonzalez-Mendez et al., 2019; Gradilla et al., 2018; Hall et al., 2019; Hall et al., 2021; Kornberg and Roy, 2014; Sanders et al., 2013). We used a double-transgenic fish 2.4Shha-ABC:Gal4FF; Shh-mChe<UAS>GFP-farn with bidirectional UAS-driven expression to visualize both Shh and plasma membranes in the MFP. Shh-mCherry was indeed detected along and at the tip of filopodia as shown at 48 hpf (Fig. 5A-A’). To test whether physiological modifications of H_2_O_2_ levels could impact filopodia, we expressed DAO at the level of the plasma membrane in MFP cells (2.4Shha:Gal4; UAS:Igk-mb5-DAO-mCherry), and we injected D-Ala into the spinal cord canal at 48 hpf. We then visualized the filopodia using the fluorescence of the membrane-bound mCherry (Fig. 5B) and quantified the number of filopodia per cell (Fig. 5C) as well as the length of the filopodia (Fig. 5D). Interestingly, enhancement of H_2_O_2_ levels induced increases in both the number and the length of filopodia (Fig. 5C-D) *in vivo*, suggesting that mild modifications of H_2_O_2_ levels could modulate Shh functioning in the neural tube. To test this hypothesis, we treated zebrafish larvae (2.4Shha:Gal4; UAS:HyPer7) with NOX-i (NADPH oxidase pan inhibitor) to reduce H_2_O_2_ levels (Supplementary Fig. S1) from 30 hpf to 48 hpf. This reduction in H_2_O_2_ levels induces a decrease in filopodia number in MFP (Supplementary Fig. S1). We then analysed the distribution of OPCs in the embryonic spinal cord at 72 hpf in (Olig2:EGFP) larvae (Fig. 5E-F). A small reduction in H_2_O_2_ levels (approximately 10%) was sufficient to enhance by a factor of 2 the generation of OPC (Fig. 5F), known to dependt on Shh activity (Al Oustah et al., 2014), without affecting *shh* expression (not shown). Thus, small modifications in H_2_O_2_ levels not only modified the number and length of filopodia *in vivo* but also altered a Shh-dependent switch.

**Fig. 5.**
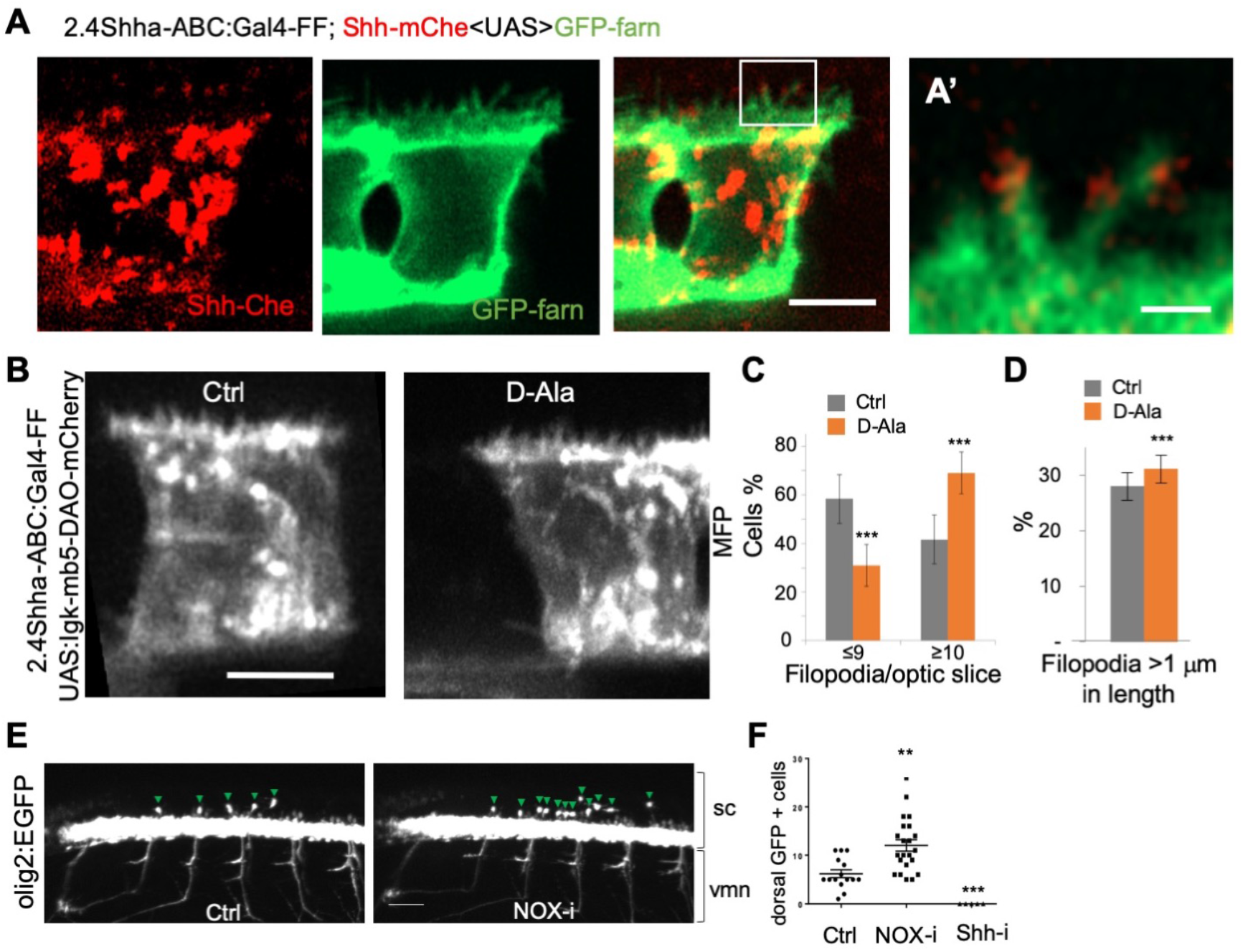
H_2_O_2_ modulates filopodia production and OPC bihaviour in the zebrafish spinal cord. (**A**) Shh-mCherry visualization in MFP filopodia in live embryos at 48 hpf (Scale bar: 5 μm). (**A’**) Enlarged view (scale bar: 1 μm). (**B**) Visualization of filopodia at 48 hpf in Tg(2.4Shha-ABC:Gal4-FF;UAS:Igk-mb5-DAO-mCherry) before or after D-Ala (10 mM) injection (scale bar: 5 μm). (**C**) Quantification of filopodia in MFP cells (see Methods) in Ctrl and D-Ala-injected larvae. (**D**) Proportion of filopodia longer than 1 μm in MFPs of Ctrl and D-Ala-injected larvae. (**E**) Detection of GFP at 72 hpf in *Tg(olig2:EGFP)* larvae incubated from 30 hpf in control solution or in NOX-i at 10 μM. (**F**) Quantification of OPCs generated at 72 hpf. Details on statistics in Methods. sc: spinal cord; vmn: ventral motor nerves.

### Shh regulates H_2_O_2_ levels and filopodial growth via a non-canonical route

After demonstrating that H_2_O_2_ impacts Shh trafficking (Fig. 3) and signaling outcome (Fig. 5), we wondered whether this was integrated into a larger H_2_O_2_-Shh feedback loop, as first observed during adult zebrafish caudal fin regeneration (Meda et al., 2016). If this were the case, Shh’s interaction with cells should itself impact H_2_O_2_ levels. We first observed that HeLa cells co-expressing HyPer and Shh-mCherry exhibited higher levels of H_2_O_2_ than cells expressing HyPer and mCherry (Fig. 6A), suggesting that Shh *per se* affected H_2_O_2_ balance. Interestingly, untransfected neighbours of cells expressing Shh-mCherry also displayed increased H_2_O_2_ levels compared to neighbours of cells expressing only mCherry (Fig. 6A). This observation suggests a paracrine effect of Shh on H_2_O_2_ levels. Moreover, when HyPer was addressed to the plasma membrane (Lck-HyPer), filopodia bearing Shh were readily identified, and high levels of H_2_O_2_ consistently co-localized with Shh (Fig. 6B). To quantitively assess the impact of Shh on H_2_O_2_ levels, we treated a stable cell line expressing Lck-HyPer with Shh-containing conditioned medium. This treatment induced a global increase in H_2_O_2_ levels (Fig. 6C-D), which was accompanied by filopodia growth, as quantified by their cumulative length (Fig. 6E). Finally, we investigated the molecular pathway used by Shh to increase H_2_O_2_ levels. Smoothened involvement was highly unlikely since its mRNA was not detected in our RT-qPCR experiments (Supplementary Table S1). Indeed, cyclopamine (Shh-i), a Shh antagonist that specifically binds Smo, did not affect basal H_2_O_2_ levels and did not inhibit the Shh-induced H_2_O_2_ burst (Fig. 6F). In contrast, treatment with Nox-i efficiently inhibited Shh-mediated H_2_O_2_ increase, demonstrating the involvement of NOX enzymes (Fig. 6F). Rac1 was an attractive candidate to bridge Shh and NOX in our system. It is ubiquitous, activates NOX2 (the only NOX complex present in HeLa cells, Supplementary Table S1), and is well known to regulate cytoskeleton dynamics (Acevedo and Gonzalez-Billault, 2018); in addition, its interplay with Shh and NOX1 was previously observed in a different system (Polizio et al., 2011). We used a Rac1 inhibitor (Rac1-i) to evaluate its involvement in H_2_O_2_ production induced by Shh treatment. As shown in Figure 6G, Rac1-i itself had no effect on H_2_O_2_ levels in untreated cells but was able to block the effect of Shh treatment. Rac1 thus sits at the crossroads of Shh activity on HeLa cells, leading both to H_2_O_2_ production via the NOX/SOD complex and to cytoskeleton remodeling that ultimately enables filopodium formation. A potential pathway to mediate Rac1 activation by Shh was recently published (Makihara et al., 2018): Shh-mediated axon guidance in the spinal cord depends on BOC receptor stimulation, leading to ELMO-Dock release and Rac1 activation. Since BOC, ELMO2 and 3, as well as several DOCK proteins, are expressed in HeLa cells (Supplementary Table S1), we tested this hypothesis using a pan-Dock inhibitor (CPYPP, Dock-i). As shown in Fig. 6G, DOCK-i *per se* did not affect H_2_O_2_ levels but blocked the H_2_O_2_ increase induced by Shh. Moreover, simultaneous treatment with DOCK-i and Rac-i had no cumulative effect (Fig. 6G), suggesting that they belong to the same pathway. In summary, these experiments show that Shh induces a local increase in H_2_O_2_ levels in HeLa cells via a non-canonical route involving its receptor Boc and the Elmo-Dock pathway to activate Rac1.

**Fig. 6:**
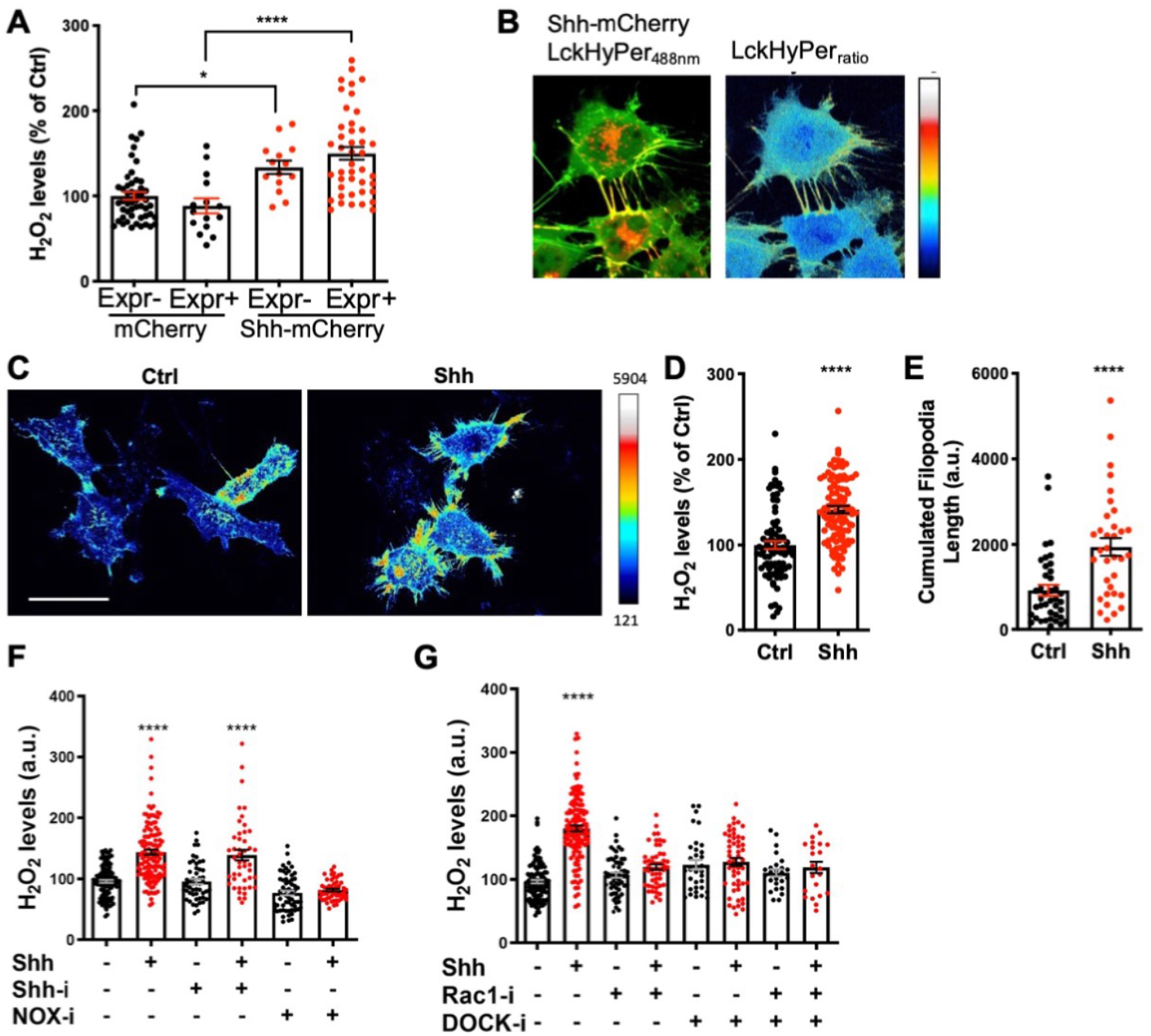
Shh regulates H_2_O_2_ levels and filopodial growth. (**A-E**) Cells stably expressing membrane-bound HyPer (**A-B**) either transfected with constructs coding for Shh-Cherry (**A, B**) or Cherry (**A**), were (**C-E**) treated (Shh) or not (Ctrl) with Shh, or were (**F, G**) treated with various inhibitors. (**A**) H_2_O_2_ was measured separately in cells expressing (Expr+) and non-expressing (Expr-) the transfected construct, coding for either Shh-mCherry (right) or Cherry (left). H_2_O_2_ levels in the non-expressing cells transfected with mCherry were taken as 100%. (**B**) Merging the red (Shh-mCherry) and green (Lck-Hyper_488nm_) signals highlights Shh-bearing filopodia (left), where H_2_O_2_ levels are distinctly elevated (right). (**C-E**) Treated cells (Shh) display higher H_2_O_2_ levels (imaged in **C**, quantified in **D**) and have longer filopodia (imaged in **C**, quantified in **E**) than untreated cells (Ctrl). (**F**) Cells treated (or not) with Shh were treated (or not) with either cyclopamine (Shh-i) or VAS2870 (NOX-i). (**G**) Cells treated (or not) with Shh were treated (or not) with either NSC23766 (Rac1-i) or CPYPP (DOCK-i) alone or in combination. In **D** and **F-G**, untreated cells were taken as 100%. Details on statistics in Methods. Scale bar: 50 μm.

As thoroughly discussed in Acevedo and Gonzalez-Billault (Acevedo and Gonzalez-Billault, 2018), this leads to NOX2 activation along the redox branch of Rac1 activity and membrane protrusion growth along the actin cytoskeleton branch. Very recently, Shh cytoneme formation in rodent cells was demonstrated to depend on another feedback loop, including a Disp-BOC co-receptor complex (Hall et al., 2021).

Signaling filopodia or cytonemes are a common route for the secretion of morphogens (Gradilla et al., 2018; Ramirez-Weber and Kornberg, 1999) and it was recently shown that several morphogens (Fgf2, Wnt8a, Wnt3a, Shh) are involved in a regulatory loop with cytoneme formation and growth, independent of their canonical pathway (Hall et al., 2021; Mattes et al., 2018). This happens during embryonic development, at a time when H_2_O_2_ levels are highly dynamic (Rampon et al., 2018).

We report here a mechanism by which physiological levels of H_2_O_2_ adjust the trafficking of Shh, which, in turn, enhances H_2_O_2_ levels. As a consequence, local and timely H_2_O_2_ production may polarize the journey of Shh by modifying the rates of Shh secretion and endocytosis as well as the regulation of filopodia. We speculate that H_2_O_2_ could integrate the spreading of morphogens in a developing tissue, with different morphogens responding differentially to specific concentrations of H_2_O_2_ and, in turn, modifying H_2_O_2_ levels. Indeed, we already know that H_2_O_2_ impact on trafficking differs between Engrailed (Amblard et al., 2020b) and Shh (this study). This interplay between morphogens and redox signaling is also likely to have a role in various pathologies (Sies, 2017).

In conclusion, we propose that environmental insults or individual genetic variations inducing very subtle differences in H_2_O_2_ levels could impact morphogen distribution, resulting in interindividual differences in organism patterning or disease susceptibility.

## Materials and methods

### Fish husbandry and pharmacological treatments

Zebrafish were maintained and staged as previously described (Amblard et al., 2020b). Experiments were performed using the standard AB wild-type strain. The embryos were incubated at 28°C. Developmental stages were determined and indicated as hours postfertilization (hpf). The animal facility obtained permission from the French Ministry of Agriculture for all the experiments described in this manuscript (agreement no. C 75-05-12). To decrease H_2_O_2_ levels, embryos were incubated in VAS-2870 (NOX-i) (100 nM) from Enzo Life Sciences (#BML-El395-0010, Enzo Life Sciences, Inc.; Farmingdale, NY, USA) or an equivalent amount of DMSO as a control for the duration of the time-lapse analysis. To enhance H_2_O_2_ levels D-Ala (Sigma-Aldrich, #A7377) (10 mM) was injected in zebrafish ventricles one hour prior H_2_O_2_ levels or filopodia analysis.

### Expression constructs, permanent cell lines, fish transgenic lines

All recombinant DNA (Supplementary Table 3) were prepared by standard cloning methods. Plasmids and sequences are available on request.

Stable cell lines (Supplementary Table 2) were prepared under Hygromycin selection using the HeLa FlpIn-TREX cell line kindly provided by Stephen Taylor (Tighe et al., 2008) expressing the tetracycline repressor (T-Rex, Life Technologies).

Transgenic lines used in this study were: *2.4Shha-ABC:Gal4* (this article); *UAS:Igkmb5DAOmCherry* (this article); *UAS:Igkmb5CATmCherry* (this article); *UAS:Shh-mCherry, GFP-farn* (this article); *UAS:HyPer7* (this article); *olig2:GFP* (Shin et al., 2003). The transgenic fish lines were constructed as described (Urasaki et al., 2006) using pTol2 derivatives containing the appropriate promoter/enhancer and the SV40 late polyadenylation signal (SVLpA). Shh:Gal4 contains the Gal4 DNA binding domain fused to 2 minimal activator sequences (Gal4BD-FF(Seipel et al., 1992)) inserted between the −2.4Shha promoter and ar-A, ar-B, ar-C Shha enhancers (a kind gift of R. Ertzer (Ertzer et al., 2007)). UAS:HyPer7 contains HyPer7 sequence (Pak et al., 2020) down 5xUAS (derived from(Akitake et al., 2011)). UAS:igkmb5-DAOmCherry and UAS:igkmb5-CATmCherry contain, downstream the same enhancer and provided with a signal peptide (Igk) from kappa light chain and a minimal transmembrane domain (mb5) from CD4, fusions of mCherry (in C-terminal position) with either D-Aminoacid oxidase (DAO(Matlashov et al., 2014)) or mouse Catalase deprived from its lysosome-targeting signal (CAT). For bidirectional expression the 5xUAS regulatory element was flanked by minimal promoter/5’ UTR sequences from pCS2 on one side and CMV on the other side (Anton et al., 2018).

### Embryo live imaging and image processing

The larvae were anesthetized in tricaine solution and embedded in low-melting agarose (0.8 %). Imaging was performed with a CSU-W1 Yokogawa spinning disk coupled to a Zeiss Axio Observer Z1 inverted microscope equipped with a sCMOS Hamamatsu camera and a 10× (Zeiss 0.45 Dry WD: 2.1mm) or a 25× (Zeiss 0.8 Imm DIC WD: 0.57 mm) oil objective. DPSS 100 mW 405 nm and 150 mW 491 nm lasers and a 525/50 bandpass emission filter were used for HyPer7 imaging, and DPSS 100mW 561 nm laser and a BP 595/50 was used for mCherry imaging. Floor plate cells imaging was performed using a Zeiss LSM 980 - AiryScan 2 confocal equipped with a AiryScan detector GaAsP-PMT and a 25× (Zeiss 0.8 Imm WD: 0.57mm) or a 40× (Zeiss 1.3 Oil DIC (UV) WD: 0.22mm) oil objectives. DPSS 10 mW 488 nm and 10 mW 561 nm lasers and respectively 517 nm and 610 nm AiryScan emission filters were used for GFP and mCherry acquisition. AiryScan SR mode was used and AiryScan-processed images were analysed. To calculate the HyPer ratio, images were treated as previously described(Mishina et al., 2013). For filopodia analysis, 48 hpf Tg(2.4Shha-ABC:Gal4-FF) larvae expressing Igkmb5-DAO-mCherry were used for fluorescence acquisition as described above, and one slice of each Zstack was extracted. Slices presenting the maximum filopodia number were selected for FiloQuant processing and analysis (Jacquemet et al., 2019).

### Pharmacological treatments

To decrease H_2_O_2_ levels, cells were treated with extracellular Catalase (Sigma-Aldrich, #C1345, 400 U/mL). To increase H_2_O_2_ levels, cells expressing D-amino acid oxidase (DAO) were treated with 10 mM D-alanine (Sigma-Aldrich, #A7377) before the internalization or secretion assays were performed. To inhibit NOX activity, cells were pre-treated for 1 h with 10 μM VAS-2870 (NOX-i; #BML-El395-0010, Enzo Life Sciences, Inc.; Farmingdale, NY, USA) or an equivalent amount of DMSO as a control. To inhibit Rac1, cells were pre-treated for 6 h with 20 μM NSC23766, a Rac1-inhibitor (Rac1-i; No2161, Tocris). To inhibit Dock release from Elmo, cells were pre-treated for 6 hours with 100 μM CPYPP (DOCK-i; No4568, Tocris). To inhibit Shh signaling, cells were pretreated for 24 h with 10 μM cyclopamine (Shh-i; #239803, Millipore).

### Quantitative secretion assay

Cells (13,000 per well) were plated on 96-well plates (Greiner Bio-one) coated with polyornithine (10 μg/mL). After 10 h, the cells were co-transfected with a plasmid expressing Shh-SBP-HiBiT (or a secreted control, SecGFP-SBP-HiBiT) under doxycycline control and a plasmid constitutively expressing the hook (core streptavidin-KDEL provided with a signal peptide). Some of these cells received an additional plasmid constitutively expressing a membrane-bound fusion between mCherry and D-Aminoacid Oxidase (Lck-Che-DAO). After 24 h medium was removed and cells were incubated with fresh medium containing doxycycline for 2 h (to induce HiBiT fusion expression). Medium was changed and secretion was induced with biotin (100 μM final) and, after purified LgBiT protein addition in the medium, luciferase activity was measured 1 h later with a 96-well plate luminometer (Tristar, Berthold) as described in the HiBiT assay kit (Promega). The cells were then lysed to measure intracellular protein expression. Normalization with biotin-untreated wells enabled us to calculate the secretion index and report the secretion efficiency.

### Quantitative internalization assay

Cells (90,000 per well) stably expressing LgBiT were plated in 24-well culture dishes. After 24 h, medium was changed and cells were incubated for 30 min with medium containing HiBiT fusions (taken from cells expressing Shh-HiBiT or SecCh-HiBiT for 48 h, and adjusted on protein Luciferase activity), before incubating the cells with trypsin and removing them. After centrifugation cells were lysed and the luciferase activity of endocytosed protein was measured with a 96-well plate luminometer (Tristar, Berthold) with a HiBiT assay kit (Promega).

### Quantitative intercellular transfer assay

Cells (90,000 per well) were plated in 24-well culture dishes. After 10 h, the cells were cotransfected with a plasmid expressing Shh-SBP-HiBiT under the control of doxycycline and a plasmid constitutively expressing Lck-Che-DAO. After 24 h, cells were removed using trypsin and co-cultured on 96-well plates (Greiner Bio-one) coated with polyornithine (10μg/ml) with cells stably expressing LgBiT anchored to the extracellular side of the cell surface (siL-LgBiT-mb5). After 5 h, cells were incubated with fresh medium containing doxycycline for 2 h (to induce Shh-SBP-HiBiT expression). Medium was then changed and secretion was induced with biotin (100 μM final), and luciferase activity was measured every 1 h over 4 h with a 96-well plate luminometer (Tristar, Berthold) as described in the HiBit assay kit (Promega).

### H_2_O_2_ imaging with the HyPer probe in HeLa cells

HyPer fluorescence was excited with DPSS 10 mW 488 nm and 10 mW 405 nm lasers, and the corresponding YFP emission was measured using a 525/50 bandpass emission filter. Spinning-disk images were acquired using a 63× objective (63×/1.4 oil WD: 0.17 mm) on a Spinning-Disk CSU-W1 (Yokogawa) equipped with a sCMOS Hamamatsu 2048×2048 camera. To calculate the HyPer ratio, images were treated as previously described (Mishina et al., 2013).

### Quantification and statistics

Data were analyzed using GraphPad Prism 6 and expressed as the mean ± standard error of the mean (SEM). Statistical significance was calculated using a 2-sided paired Student’s *t*-test. For multiple conditions, ordinary one-way ANOVA followed by Tukey’s multiple comparison test was performed to evaluate the significant differences. For filopodia analysis, statistical errors (SD) were estimated as √p(1-p)/n, where p is the percentage in a class and n is the total number investigated (or √1/n when p=0 or 1).The degree of significance was represented as follows: * p-value ≤ 0.05; ** p-value ≤ 0.01; *** p-value ≤ 0.001; and **** p-value ≤ 0.0001. Sample sizes and number of replicates are given in Supplementary Table 4.

## Details of author contributions

AJ, MV and SV conceived the project and designed the experiments. IA, AJ, IQ, MT, and MV prepared the DNA constructs, cell lines and zebrafish lines used in this study. IA, AJ, AM, CR, MT and SV performed the experiments. AG and CL provided the HMBR and useful advices. IA, CR, MT, MV and SV analyzed the experimental data. IA, MV and SV wrote the paper with comments from all authors.

## Acknowledgements and funding sources

We thank Helmut Sies (Heinrich Heine University Düsseldorf, D) for helpful discussions, Sylvie Schneider-Maunoury (Sorbone Université, F) and Christine Vesque (Sorbone Université, F) for fruitful comments on the manuscript. This work was performed at the Collège de France and benefited from Alain Prochiantz’s constant support. The authors are grateful to Stephen Taylor (University of Manchester, UK) for providing the HeLa Flp-In cell and to Ariel Ruiz i Altaba (Université de Genève, CH) for providing mouse Shh and mouse Shh::GFP expressing plasmids, the second one being derived from an original construct from Andrew McMahon’s laboratory (Harvard, USA). This work received support under the program “Investissements d’Avenir” launched by the French Government and implemented by the ANR with the following references: ANR-10-LABX-54 MEMO LIFE - ANR-11-IDEX-0001-02 PSL* Research University.

