## supplementary for "Reciprocal regulation of Shh trafficking and H_2_O_2_ levels via a noncanonical BOC-Rac1 pathway"

**Contents**

**Figure S1**
**Tables S1-S4**

**Figure S1. NOX-I treatment induces a reduction of $H_2O_2$ levels as well as a reduction of filopodia length in the MFP**.

**A**, $H_2O_2$ levels decreased after NOX-i treatment. 2.4Shha-ABC:GalA-FF/UAS:HyPer7 embryos were incubated in NOX-i (10 μM), and $H_2O_2$ levels were quantified in the MFP at 46 hpf. Representative images are shown. fp: floor plate; nc: notochord. Scale Bar: 10 μm.

**B,** Quantification of filopodia length in MFP cells (see Methods) in Ctrl and NOX-I treated larvae.

Details on statistics in Material and Methods

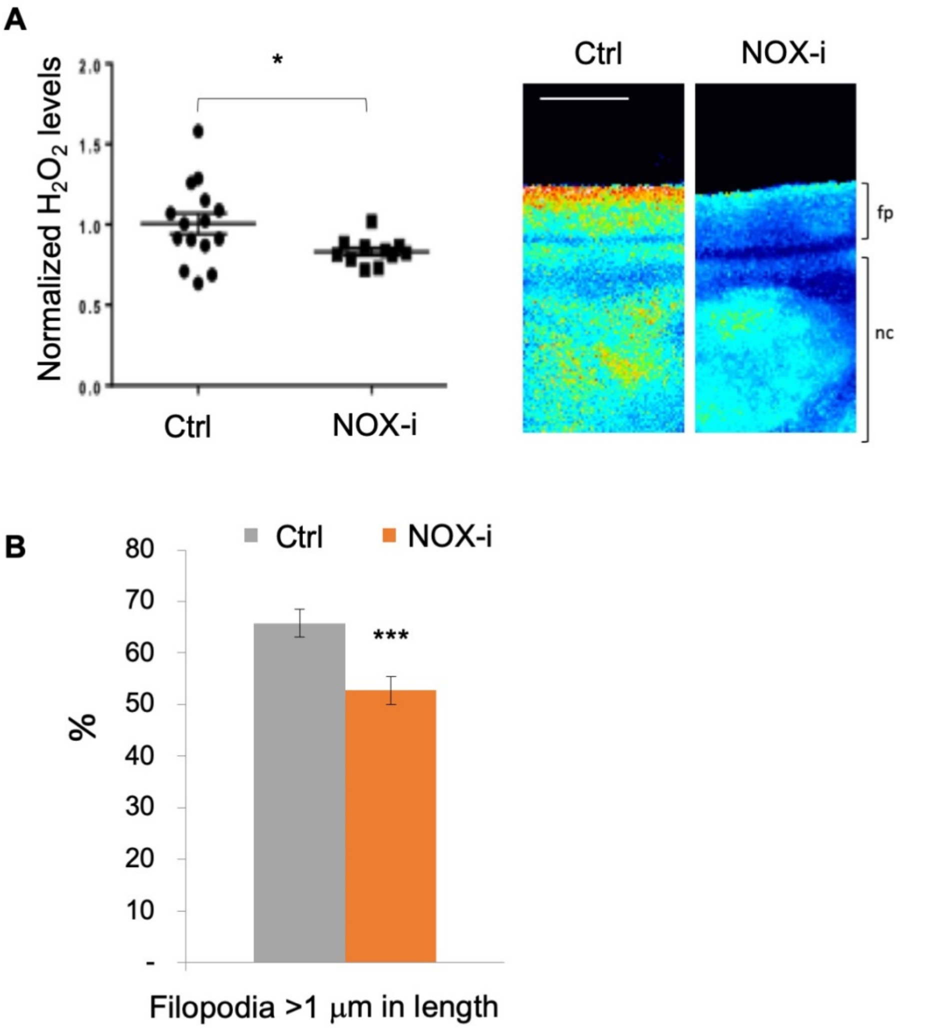

**Table S1**

nd : non detected, un : untested

positive value indicates that gene of interest is x fold more expressed than gapdh and negative values x fold less than gapdh

| | Hela ΔCt | HEK | Universal probe ID | Left primer | Right primer |
|---|---|---|---|---|---|
| ***gadph*** | | | 68 | ccc cgg ttt cta taa att gag c | ctt ccc cat ggt gtc tga g |
| ***hhat*** | 12.6±2.2 | un | 22 | gct ggg agt cac tgt gga g | gct tgt ggg gag aag tat cg |
| ***disp1*** | 7.0±1.4 | un | 43 | tgt gca atg tag ata att cca gga t | gcg atg tag ttt ccc agt gtc |
| ***ptch1*** | 3.8±0.2 | un | 17 | tgg gat taa aag cag cga ac | cga ctt act cgt cct cca act t |
| ***ptch2*** | -4.8±1.1 | un | 17 | tgg aac agc tct ggg tag aag | ccc agc ttc tcc ttg gtg ta |
| ***scube2*** | -45±9.3 | un | 17 | agc tgc cat cca cag tac aa | cca ctg atg tgg tgt tgc tc |
| ***smo*** | nd | -3.2 | 22 | gct tcc ggg act atg tgc ta | aaa cat ggc aaa cag gtt ga |
| ***boc*** | 3.3±4.6 | un | 6 | cgc caa ccc tct act atg tgg t | aat gcc aga gat ggt cca at |

| | | | | | |
|---|---|---|---|---|---|
| ***nox1*** | nd | un | 43 | ccc agc aga agg ttg tga tt | ttg agg ggc aat taa caa aga |
| ***nox2*** | -5.2±0.5 | un | 6 | aca att gca agt caa cac cct a | ctc agg gtt tca gcc aag g |
| ***nox3*** | nd | un | 22 | tca agt cca ggc att gtg tt | aac agg ccg tca cag gaa |
| ***nox4*** | nd | un | 22 | aac caa ggg cca gag tat ca | gct gag gct ctg ctt aga cac |

**Table S2: Stable HeLa cell lines used in this study**

| Name | Construct used for HeLa Flp-In | Expressed protein | ref |
|---|---|---|---|
| GBi | pcDNA7LGBi | LgBiT | this study |
| siLGBimb5 | pcDNA4siLGBiCD5 | IL2-LgBiT-mb5 | this study |
| LckHyPer | pcDNA7LckHyPer | Lck-HyPer1 | this study |
| HyPer | pcDNA7Hyper | HyPer1 | [23] |

**Table S3: Plasmids used in this study**

| Plasmid number | Plasmid name | Regulatory sequences | Expressed protein | purpose | ref |
|---|---|---|---|---|---|
| 65 | pCS2transposase | SP6 | Tol2 transposase | transgenesis | [53] |
| 1070 | pT22i2Shh24Gal4F | 2.4Shha-ABC | Gal4BD-FF | transgenesis | this paper |
| 1190 | pT22i5uasHyper7 | 5xUAS | HyPer7 | transgenesis | this paper |
| 1242 | pT22i5uasm4C5DaoChe | 5xUAS | Igk-mb5-DAO-mCherry | transgenesis | this paper |
| 1243 | pT22i5uasm4C5CatChe | 5xUAS | Igk-mb5-CAT-mCherry | transgenesis | this paper |
| 1261 | pT2i5biGfpF•Shh-SbpChe | 5xUAS bidirectional | (i) GFP-Farn<br>(ii) Shh-SBP-mCherry | transgenesis | this paper |
| 237 | pCSmChF | SP6 | mCherry-Farn | mRNA synthesis | this paper |
| 1313 | pcDNA7LGBi | CMV-tetOx2 | LgBiT | stable cell line | [43] |
| 1331 | pcDNA4LsiLGBiCD5 | CMV | IL2-LgBiT-mb5 | stable cell line | this paper |
| 754 | pcDNA7Hyper1 | CMV-tetOx2 | HyPer1 | stable cell line | [23] |
| 1036 | pcDNA2LckHyper | CMV-tetOx2 | Lck-HyPer1 | stable cell line | this paper |
| 911 | pT2iC6mShhYfaChe | sCMV | Shh-YFast-mCherry | transient *ex vivo* expr | this paper |
| 669 | pT2iC6mShhChe | sCMV | Shh-mCherry | transient *ex vivo* expr | this paper |
| 1330 | pKhCTagScLgBiT | T7 | CTagSc-LgBiT | bac prod | this paper |
| 832 | pStrKdel_ShhSbpChe | CMV | sti-STRP-kdel-ires-Shh-SBP-mCherry | transient *ex vivo* expr | this paper |
| 1265 | pCL9StKDEL | CMV | sti-STRP-kdel | transient *ex vivo* expr | this paper |
| 1237 | pcDNA2SbpShhSbi | CMV-tetOx2 | Shh-Sbp-HiBiT | transient *ex vivo* expr | this paper |
| 1311 | pcDNA2SilGfpSbpSbi | CMV-tetOx2 | IL2-GFP-SBP-HiBiT | transient *ex vivo* expr | this paper |
| 1312 | pcDNA2silSbiChe | CMV-tetOx2 | IL2-HiBiT-Che | transient *ex vivo* expr | this paper |
| 966 | pcDNA2LckCheDao | CMV-tetOx2 | Lck-mCherry-DAO | transient *ex vivo* expr | this paper |
| 669 | pT2iC6mShhChe | sCMV | Shh-mCherry | transient *ex vivo* expr | this paper |

**Table S4**
Sample sizes and number of replicates.

| Figure | condition | Sample size | Mean | Std. Deviation | Std. Error of Mean | P-value | Confidence interval |
|---|---|---|---|---|---|---|---|
| **Figure 2** | | | | | | | |
| 2C | Shh | 4 | | | | | |
| | SecGFP | 4 | | | | | |
| 2E | Shh t=5min | 2 | 16 | 1.697 | 1.2 | T=5min<br>Shh vs SecmCh<br><br>t=30min<br>Shh vs SecmCh<br>P<0.0001<br><br>t=60min<br>Shh vs SecmCh<br>P<0.0001<br><br>t=75min<br>Shh vs SecmCh<br>P=0.0491 | |
| | Shh t=30min | 8 | 100 | 23.11 | 8.17 | | |
| | Shh t=60min | 5 | 104.2 | 16.62 | 7.432 | | |
| | Shh t=75min | 2 | 112 | 5.657 | 4 | | |
| | Secm Ch t=5min | 2 | 11.5 | 7.778 | 5.5 | | |
| | SecmCh t=30min | 7 | 18.43 | 16.22 | 6.129 | | |
| | SecmCh t=60min | 4 | 36 | 7.348 | 3.674 | | |
| | SecmCh t=75min | 2 | 25.5 | 27.58 | 19.5 | | |
| 2G | Ctrl (A) | 11 | 0.1 | 0.02246 | 0.006772 | A vs B p>0.9999<br>A vs C p=0.7699<br>A vs D p=0.0919<br>A vs E p<0.0001<br>A vs F p<0.0001 | |
| | 1h (B) | 10 | 0.1017 | 0.03336 | 0.01055 | | |
| | 2h (C) | 6 | 0.1387 | 0.0549 | 0.02241 | | |
| | 3h (D) | 8 | 0.1726 | 0.05061 | 0.01789 | | |
| | 4h (E) | 7 | 0.2578 | 0.07452 | 0.02816 | | |
| | 4h30 (F) | 8 | 0.2518 | 0.09564 | 0.03381 | | |
| | Nox-i | 11 | 0.83 | 0,08184 | 0,02467 | | |

| | | | | | | |
|---|---|---|---|---|---|---|
| **Figure 3B** | | | | | | |
| Shh | Ctrl | 14 | 100 | 25.46 | 6.804 | 0.0002 |
| | D-Ala | 13 | 53.96 | 25.69 | 7.126 | |
| SecGFP | Ctrl | 14 | 100 | 30.19 | 8.069 | 0.9990 |
| | D-Ala | 12 | 101.7 | 22.06 | 6.368 | |
| **Figure 3C** | | | | | | |
| Shh | Ctrl | 15 | 100 | 16.06 | 4.147 | <0.0001 |
| | CAT | 13 | 155.5 | 23.20 | 6.434 | |
| SecGFP | Ctrl | 8 | 100 | 9.794 | 3.463 | 0.0781 |
| | CAT | 7 | 75.71 | 21.90 | 8.277 | |
| **Figure 3E** | | | | | | |
| Shh | Ctrl | 8 | 100 | 24.91 | 8.808 | 0.0082 |
| | D-Ala | 8 | 144.6 | 39.1 | 13.83 | |
| SecCh | Ctrl | 8 | 7.579 | 21.88 | 7.736 | 0.9884 |
| | D-Ala | 7 | 12 | 11.45 | 4.326 | |
| **Figure 3F** | | | | | | |
| Shh | Ctrl | 6 | 100 | 19.18 | 7.832 | 0.0017 |
| | CAT | 6 | 52.46 | 8.507 | 3.473 | |
| SecCh | Ctrl | 5 | 32.7 | 23.72 | 10.61 | 0.9869 |
| | CAT | 5 | 28.78 | 21.23 | 9.493 | |
| **Figure 3H** | | | | | | |

| | | | | | | |
|---|---|---|---|---|---|---|
| | Ctrl t=0 (A) | 10 | 0,1 | 0.0255 | 0.008065 | |
| | Ctrl t=4h (B) | 9 | 0.1395 | 0.02436 | 0.008119 | A vs B p=0.0091<br>B vs C p<0.0001 |
| | D-Ala t=4h (C) | 11 | 0.2162 | 0.03876 | 0.01169 | |
| **Figure 3I** | | | | | | |
| | Ctrl t=0 (A) | 6 | 0,1 | 0.03096 | 0.01264 | |
| | Ctrl t=4h (B) | 6 | 0.2705 | 0.07288 | 0.02975 | A vs B p<0.0001<br>B vs C p=0.0066 |
| | CAT t=4h (C) | 6 | 0.1697 | 0.02609 | 0.01065 | |

| | | | | | | | |
|---|---|---|---|---|---|---|---|
| **Figure 4B** | | | | | | | |
| | 26 hpf | 8 | 1,125 | 0,08582 | 0,03034 | <0,0001 | 0,95 |
| | 28 hpf | 8 | 1,053 | 0,07437 | 0,02629 | <0,0001 | 0,95 |
| | 30 hpf | 13 | 1,000 | 0,07127 | 0,01977 | | 0,95 |
| | 33 hpf | 5 | 0,9074 | 0,07615 | 0,03406 | <0,0001 | 0,95 |
| | 39 hpf | 5 | 0,8569 | 0,04880 | 0,02182 | 0,0005 | 0,95 |
| | 45 hpf | 5 | 0,8545 | 0,03733 | 0,01669 | 0,0027 | 0,95 |
| **Figure 4D** | | | | | | | |
| | 25 hpf | 12 | | | | | |
| | 29 hpf | 12 | | | | | |
| | 31 hpf | 24 | | | | | |
| | 35 hpf | 12 | | | | | |
| | 45 hpf | 11 | | | | | |

| | | | | | | | |
|---|---|---|---|---|---|---|---|
| **Figure 5C** | | | | | | | |
| | Ctrl ≤9 | 29 | 58.3% | 10.06 | | | |
| | Ctrl ≥10 | 29 | 41.7% | 10.06 | | | |
| | D-Ala | 24 | 31% | 8.59 | | <0.0001 | 0.95 |
| | D-Ala | 24 | 69% | 8.59 | | <0.0001 | 0.95 |
| **Figure 5D** | | | | | | | |
| | Ctrl | 243 | 20% | 2.57 | | <0.0001 | 0.95 |
| | D-Ala | 302 | 24% | 2.46 | | | |
| **Figure 5F** | | | | | | | |
| | Ctrl | 15 | 6.2 | 3.14 | 0.81 | | |
| | Nox-i | 22 | 12.05 | 5.62 | 1.2 | <0.01 | 0,95 |
| | HH-i | 5 | 0 | 0 | 0 | <0.05 | 0,95 |

| | | | | | | |
|---|---|---|---|---|---|---|
| **Figure 6A** | | | | | | |
| mCherry | Exp- (A) | 50 | 100 | 32.76 | 4.634 | A vs B p=0.757<br>C vs D p=0.5301<br>A vs C p=0.0332<br>B vs D p<0.0001 |
| | Exp+ (B) | 15 | 88.43 | 34.2 | 8.830 | |
| Shh-mCherry | Exp- (C) | 14 | 133.3 | 29.17 | 7.797 | |
| | Exp+ (D) | 43 | 150 | 50.69 | 7.716 | |
| **Figure 6D** | | | | | | |
| | Ctrl | 82 | 100 | 41.8 | 4.616 | <0.0001 |
| | Shh | 104 | 141.4 | 42.63 | 4.181 | |
| **Figure 6E** | | | | | | |

| | | | | | | |
|---|---|---|---|---|---|---|
| | Ctrl | 40 | 923.9 | 806.9 | 127.6 | <0.0001 |
| | Shh | 34 | 1940 | 1219 | 209.1 | |
| **Figure 6F** | | | | | | |
| | Ctrl (A) | 113 | 100 | 26.1 | 2.456 | A vs B p<0.0001<br>C vs D p<0.0001<br>E vs F p=0.9878 |
| | Shh (B) | 128 | 143.7 | 48.08 | 4.250 | |
| | Shh-i (C) | 50 | 95.57 | 31.02 | 4.386 | |
| | Shh+Shhi (D) | 46 | 138.9 | 57.57 | 8.488 | |
| | NOX-i (E) | 58 | 76.87 | 29.65 | 3.894 | |
| | Shh+NOX-i (F) | 54 | 81.45 | 16.33 | 2.222 | |
| **Figure 6G** | | | | | | |
| | Ctrl (A) | 111 | 100 | 29.47 | 2.798 | A vs B p<0.0001<br>C vs D p=0.7075<br>E vs F p=0.9996<br>G vs H p=0.9965 |
| | Shh (B) | 126 | 180 | 51.64 | 4.601 | |
| | Rac1-i (C) | 57 | 106.7 | 30.53 | 4.044 | |
| | Shh+Rac1-i (D) | 51 | 119.5 | 31.6 | 4.424 | |
| | DOCK-i (E) | 33 | 122.9 | 43.48 | 7.57 | |
| | Shh+DOCK-i (F) | 52 | 127.4 | 44.69 | 6.198 | |
| | Rac1-i+DOCK-i (G) | 25 | 110.3 | 28.44 | 5.688 | |
| | Shh+ Rac1-i+DOCK-I (H) | 22 | 118.7 | 41.37 | 8.821 | |

| | | | | | | | |
|---|---|---|---|---|---|---|---|
| **Figure Sup1A** | | | | | | | |
| | Ctrl | 14 | 1.0 | 0,2506 | 0,06472 | 0,030 | 0,95 |
| | NOX-i | 11 | 0.83 | 0,08184 | 0,02467 | | |

| **Figure Sup1B** | | | | | | | |
|---|---|---|---|---|---|---|---|
| | Ctrl | 298 | 65.77 | 2.75 | 0.1592 | <0.0001 | 0.95 |
| | NOX-i | 334 | 52.69 | 2.73 | 0.1495 | | |
